# Machine Learning-based Predictions of Spatial Metabolic Profiles Demonstrate the Impact of Morphology on Astrocytic Energy Metabolism

**DOI:** 10.1101/2024.09.18.613725

**Authors:** Paris Papavasileiou, Sofia Farina, Eleni D. Koronaki, Andreas G. Boudouvis, Stéphane P.A. Bordas, Alexander Skupin

## Abstract

This work introduces a machine learning framework that allows the investigation of the influence of reaction centers on the metabolic state of astrocyte cells. The proposed ML framework takes advantage of spatial astrocyte metabolic data stemming from numerical simulations for different reaction center configurations and allows for the following: (i) Discovery of cell groups of similar metabolic states and investigation of the reaction center configuration within each group. This approach allows for an analysis of the importance of the specific location of the reaction centers for a potentially critical metabolic state of the cell. (ii) Qualitative prediction of the energetic state of the cell (based on [ATP]: [ADP]) and quantitative prediction of the metabolic state of the cell by predicting the spatial average concentration of the metabolites or the complete spatial metabolic profile within the cell. (iii) Finally, the framework allows for the post hoc analysis of the developed quantitative predictive models using a SHAP approach to investigate the influence of the reaction center positions for further support of the insights drawn in steps (i)-(iii). Following the implementation of the framework, we observe that a uniform mitochondrial distribution within the cell results in the most robust energetic cell state. On the contrary, realizations of polarized mitochondrial distributions exhibit the worst overall cell health. Furthermore, we can make accurate qualitative predictions regarding cell health (*accuracy* = 0.9515, *recall* = 0.9753) and satisfactory predictions for the spatial average concentration and spatial concentration profiles of most of the metabolites involved. The techniques proposed in this study are not restricted to the dataset used. They can be easily used in other datasets that include findings from various metabolic computational models.

## Introduction

Understanding the complex interplay of molecules within cells is crucial for advancing fields such as medicine, biotechnology, and pharmacology. In the intricate landscape of cellular biology, metabolism is a complex series of interconnected pathways occurring in living cells. It operates through specific biochemical reactions and produces energy and other essential biochemical compounds. Energy in the form of ATP is the fuel of all living systems, and metabolism is designed to optimally regulate it. Metabolism is thus a prerequisite for the optimal function and survival of cells and, in extension, for the survival of organisms. An example of particularly important cells are astrocytes (cf.Fig. 1), the most abundant glial cells and crucial energetic supporters of the energy-intensive brain (***Pellerin et al., 1998***). In general, the study of cellular metabolism has evolved significantly, with researchers now employing a multidisciplinary approach that integrates both biological experiments and computational modeling. Given the intricate nature of metabolism, employing mathematical models is essential for a comprehensive investigation (***Kitano, 2002***).

**Figure 1.**
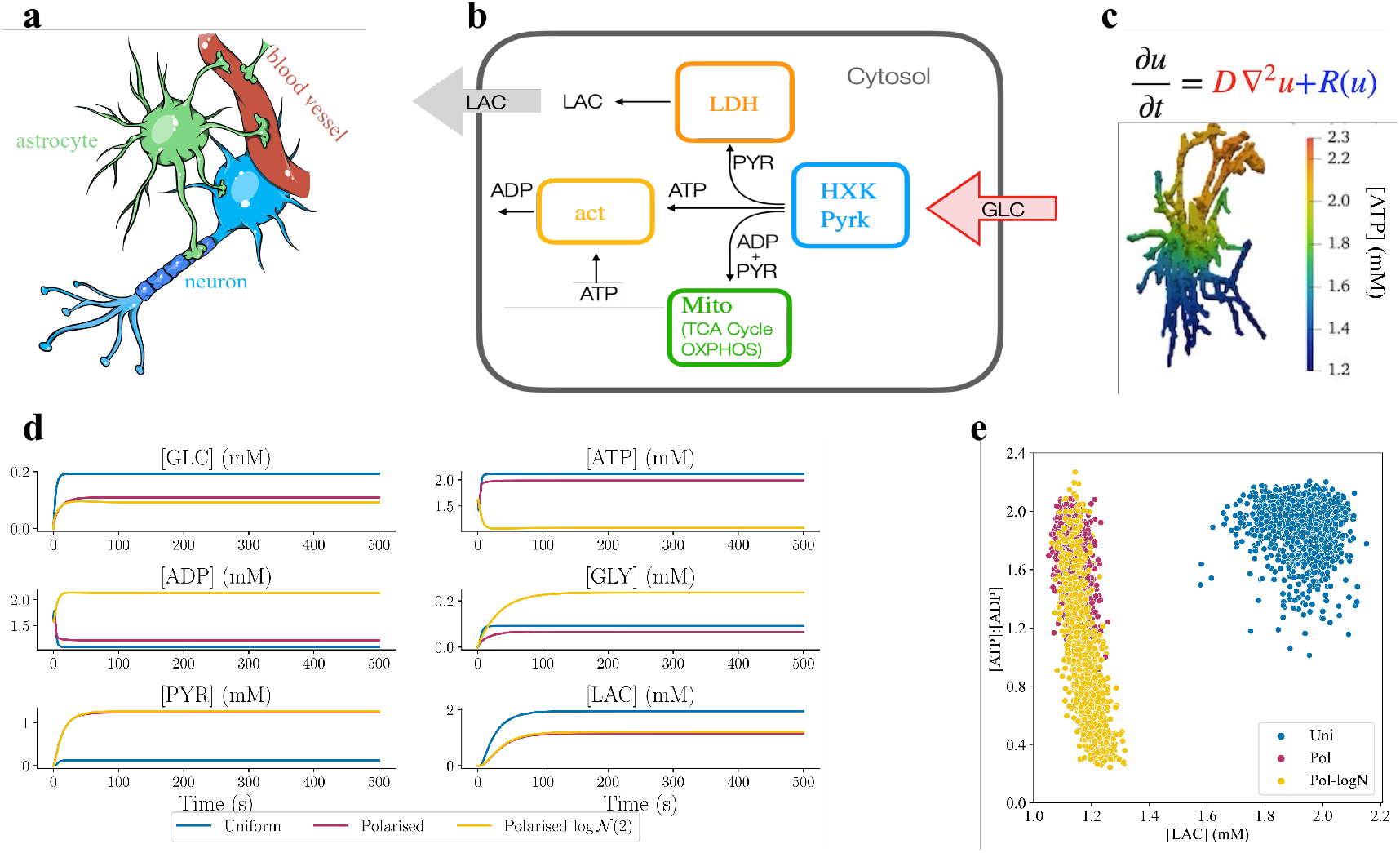
(a) A sketch illustrating the crucial location of an astrocyte between a neuron and a blood vessel, relevant for their metabolism. (b) A concise overview of the metabolic model used to describe the main pathways in astrocytes. (c) An example of [ATP] concentration profile obtained solving the metabolic model in a 3D human astrocyte obtained from a confocal microscopy image. (d) Dynamic evolution of the six considered metabolites averaged inside the 2D domain for three sampled realizations, one for each distribution of reaction centers. (e) Scatter plot of the spatial average [ATP] ∶ [ADP] vs spatial average [LAC] for all available cell configurations at steady state -Clear distinctions between uniform configurations and polarized and polarized log-normal configurations.

Metabolic processes are not uniformly distributed throughout the cell. Subcellular compartments such as mitochondria, endoplasmic reticulum, and cytoplasm exhibit distinct metabolic activities. In astrocytes, the enzymatic distribution of hexokinase seems to be fundamental for glucose uptake (***Sánchez-Alvarez et al., 2004***), while the location of mitochondria appears to be crucial for calcium activity (***Jackson and Robinson, 2015***). Thus, the current research direction aims to include spatial cellular information (***Agapakis et al., 2012***) to spatially quantify metabolites and their dynamics over time. Recent advances in analytical techniques have contributed to obtaining a snapshot of the cellular status. For example, spatial metabolomics (***Alexandrov, 2023***) aims to identify and analyze metabolites directly within their -usually-geometrically complex spatial surroundings. Imaging and image analysis techniques have been proven to be useful in investigating spatio-temporal intracellular ATP and cellular morphological changes (***Tantama and Yellen, 2014***; ***Suzuki et al., 2015***). In addition, these spatially resolved data contribute to a more comprehensive understanding of how metabolic processes are compartmentalized and coordinated within the cell.

Complementing the analytical techniques mentioned above, computational approaches have become a valuable tool for unraveling the complexity of cellular metabolism. Classical metabolic modeling approaches range from stoichiometric models (***Llaneras and Picó, 2008***) to kinetic simulations (***Cortassa et al., 2003***; ***Aubert and Costalat, 2005***; ***Aubert et al., 2005***). These models are capable of predicting and simulating the dynamics of the metabolic system. In addition, they can help guide experimental design and generate hypotheses. The main limitation of these models is the assumption of a well-mixed cellular environment, which neglects the spatial heterogeneity present in the biological systems that we discussed above. Several recent computational models have proposed spatially resolved kinetic models and agent-based simulations (***Szabó and Merks, 2013***; ***Cleri, 2019***) exploring how metabolite concentrations can vary in spatial dimension in different cellular morphologies of varying geometric complexity. This modeling approach is particularly valuable as it approaches biological reality and is well suited for the study of phenomena such as organelle crosstalk and the impact of spatial constraints on metabolic fluxes (***Khalid et al., 2018***; ***Bell et al., 2019***; ***Ellingsrud et al., 2020***; ***Farina et al., 2023***; ***Garcia et al., 2023***). Biological snapshots of metabolite concentrations obtained from *in vivo* and *in vitro* cells offer valuable glimpses into cellular states (***Kobayashi et al., 1999***) and can be used as starting points for spatially resolved models. Although these data can characterize the cellular state at the moment they are collected, they are unable to capture the dynamic nature of cellular metabolism. Moreover, there is a limit to the data that can be collected from a cellular sample: staining a cell to gain information on one metabolite can prevent the investigation of another. Lastly, the lack of comprehensive data on the temporal aspects of metabolic processes hinders the ability of computational models to accurately simulate and predict the real-time behavior of cellular metabolism. Bridging these gaps in both experimental snapshots and computational modeling data is essential for understanding the intricate dynamics that govern cellular metabolic networks.

Addressing the limitations in our current understanding of cellular metabolism, machine learning techniques (***Zampieri et al., 2019***; ***Alber et al., 2019***) could be applied to bridge the gap between static biological snapshots and dynamic models. For this purpose, we implement a machine learning approach on a dataset consisting of the results of a spatially resolved computational metabolic model of an astrocyte (***Farina et al., 2023***). This computational model provides us with spatial information of the metabolites in a simplified two-dimensional rectangular cellular domain, given different configurations of reaction centers in the form of coordinates on the *x*- and *y*-axes.

The proposed approach aims to discover reaction centers (inputs) that are potentially critical to the metabolic state of the cell (output). To this end, the following steps are necessary: a) Discovery of groups of similar metabolic profiles, using *only* the output of the computational model (spatial metabolic concentrations at steady state). This is achieved through the use of clustering algorithms. By analyzing the inputs corresponding to the resulting clusters, we can draw insights into the relationship between the reaction center position and the metabolic state of the cell. b) Qualitative prediction of cell health status using *only* the coordinates of the reaction centers as input. This is made possible through the use of classification algorithms. The input-output relationship insights derived from the previous step are expected to greatly influence the predictions of the classification algorithm used. c) Quantitative prediction of the metabolic state of the cell using the coordinates of the reaction centers as inputs. This is enabled by the use of regression algorithms, in our case, Artificial Neural Networks (ANNs). We are able to predict both the average concentration of the metabolites in the domain and the spatial profile of the metabolites in the domain at steady state. Last but not least, d) since the explainability of the developed “black-box” ANN models is very important for our application, a SHAP analysis (***Lundberg and Lee, 2017***) can be performed for the developed regression models. The obtained values can shed more light on the effect of each input (reaction center coordinates) on model output (spatial metabolite concentration) and further indicate whether the insights derived in the previous steps are meaningful.

These techniques have the potential to decode the complexity inherent in cellular metabolism, offering a means to generate more comprehensive and accurate representations of metabolic processes. Hopefully, they can also be applied to spatially resolved computational models of higher metabolic or geometric complexity, which are predominant when it comes to cells.

## Methods

### Computational Model

#### Biochemical Reaction Model

We consider a spatially resolved metabolic model, proposed in (***Farina et al., 2021, 2023***), which prioritizes the arrangement of the reaction sites in the domain and the geometries of the domain at the expense of a more elementary chemical model.

In its simplicity, the model captures the main fundamental metabolic energy pathways in five chemical reactions: glycolysis, mitochondrial activity, and lactate dehydrogenase. Glycolysis is described by two chemical reactions named HXK and PYRK. The first one accounts for the enzymes: hexokinase, phosphoglucose isomerase, phosphofructose kinase, and fructose bisphosphate aldolase. HXK consumes glucose (GLC) and adenosine triphosphate (ATP) producing adenosine diphosphate (ADP) and glyceraldehyde (GLY). The second reaction, PYRK, uses the product of the first reaction to produce ATP and pyruvate PYR. Now, PYR can either be used by the lactate dehydrogenase enzyme (LDH) to produce lactate (LAC) or enter the mitochondria and contribute to mitochondrial activity. Mitochondrial activity accounts for the Krebs cycle and oxidative phosphorylation, producing ATP. Finally, the energetic production within the cell is balanced by considering the cellular activity that consumes ATP.

The chemical model is then described as follows:

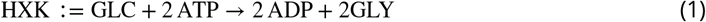

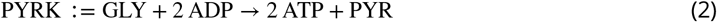

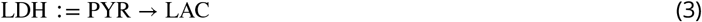

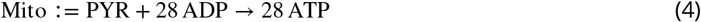

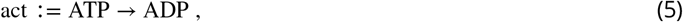

An overview of the model is presented in Fig. 1.

#### Mathematical model

Mathematically, the model is then translated into a reaction-diffusion system (***Murray, 2002***) through a set of partial differential equations (PDE), which allows us to a) solve the metabolic model in a geometrical bounded domain; b) account for the molecules’ diffusivity; c) distribute spatially the chemical reaction sites inside the domain.

In a bounded 2-dimensional domain, Ω, we consider a fixed number, *M* ∈ ℝ^+^, of reaction sites for chemical reactions: HXK, PYRK, LDH and Mito, which are spatially distributed using a spatial reaction rate density, *K*_j_ Spatial reaction rates are defined as the product between classical reaction rates *K*_*j*_, and Gaussian functions defined with a center 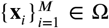 and variance σ_*i*_ ∈ ℝ^+^. The cellular activity, act, operates homogeneously in the domain Ω with reaction rate *K*_act_ .

The reaction-diffusion system is defined as follows:

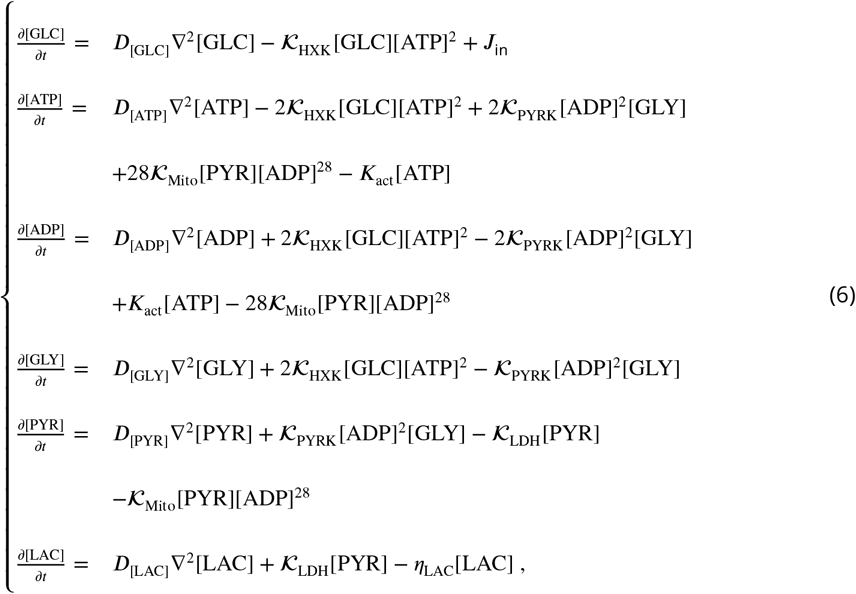

where:

- The source of GLC is described through a function *J*_in_ ∶ Ω × [0, *T* ] → ℝ:

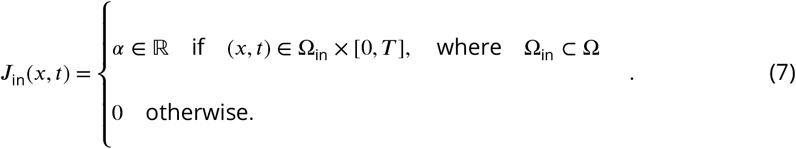
- The degradation of LAC, which is proportional to the amount of LAC in region Ω_out_ ⊂ Ω is described by the function *η*_LAC_ ∶ Ω × [0, *T* ] → ℝ

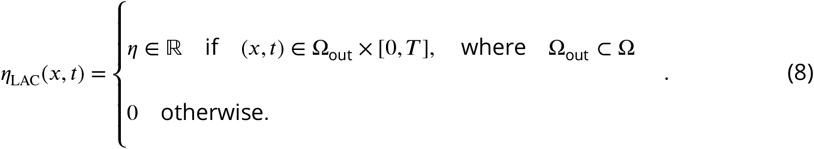

For more details on the mathematical model and parameters, we refer the readers to (***Farina et al., 2023***).

### Data acquisition

Data were acquired by numerically solving the reaction-diffusion system that arose from the metabolic model presented in the previous section. We used standard finite element methods (***Hughes, 2012***) using the FEniCS software (***Alnæs et al., 2015***). First, the reaction-diffusion system was converted to its corresponding weak form. Then, we spatially discretize the two-dimensional rectangular domain by finite elements using the package *mshr* (number of finite elements 25298 and number of dofs 13207). We temporally discretize the time derivative using a backward Euler scheme with a time step of 0.15 (s) (***Quarteroni and Valli, 2008***). The solution of the weak form is defined on the space of piecewise Lagrangian finite elements of degree one.

We consider a 2-dimensional rectangular domain ([0, *l*] × [0, *L*], with width *l* = 4 *μ*m and length *L* = 140 *μ*m) where we place 10 reaction sites per chemical reaction with a spatial extent of σ = *μ*m. The input/inlet of the system is the entrance of GLC in the bottom left corner, while the output/outlet is the outflux of LAC in the opposite corner. To investigate the crucial role of spatial arrangement in cellular domains, we consider three possible distributions of the reaction sites: uniform, polarized, and polarized log-normal. The uniform distribution considers the 10 reaction sites per chemical reaction to be sorted from uniform distributions. The polarized consider an extreme reaction site configuration supposing that glycolysis is located at the bottom of the rectangular domain close to the GLC influx while the 10 reaction sites for LDH are sorted at the top of the rectangular domain. The main difference between polarized and polarized log-normal lies in the distribution of the mitochondria. In the first case, six reaction sites for Mito are placed where glycolysis is located, and four reaction sites are at the top of a part of the rectangular domain, to ensure that some mitochondria can be found throughout the domain. The polarized log-normal setting uses a log-normal distribution to sort the 10 Mito reaction sites, causing mitochondria to be located mainly in the lower part of the domain and almost none co-located with LDH. Examples of the three distributions can be seen in Figure 2.

**Figure 2.**
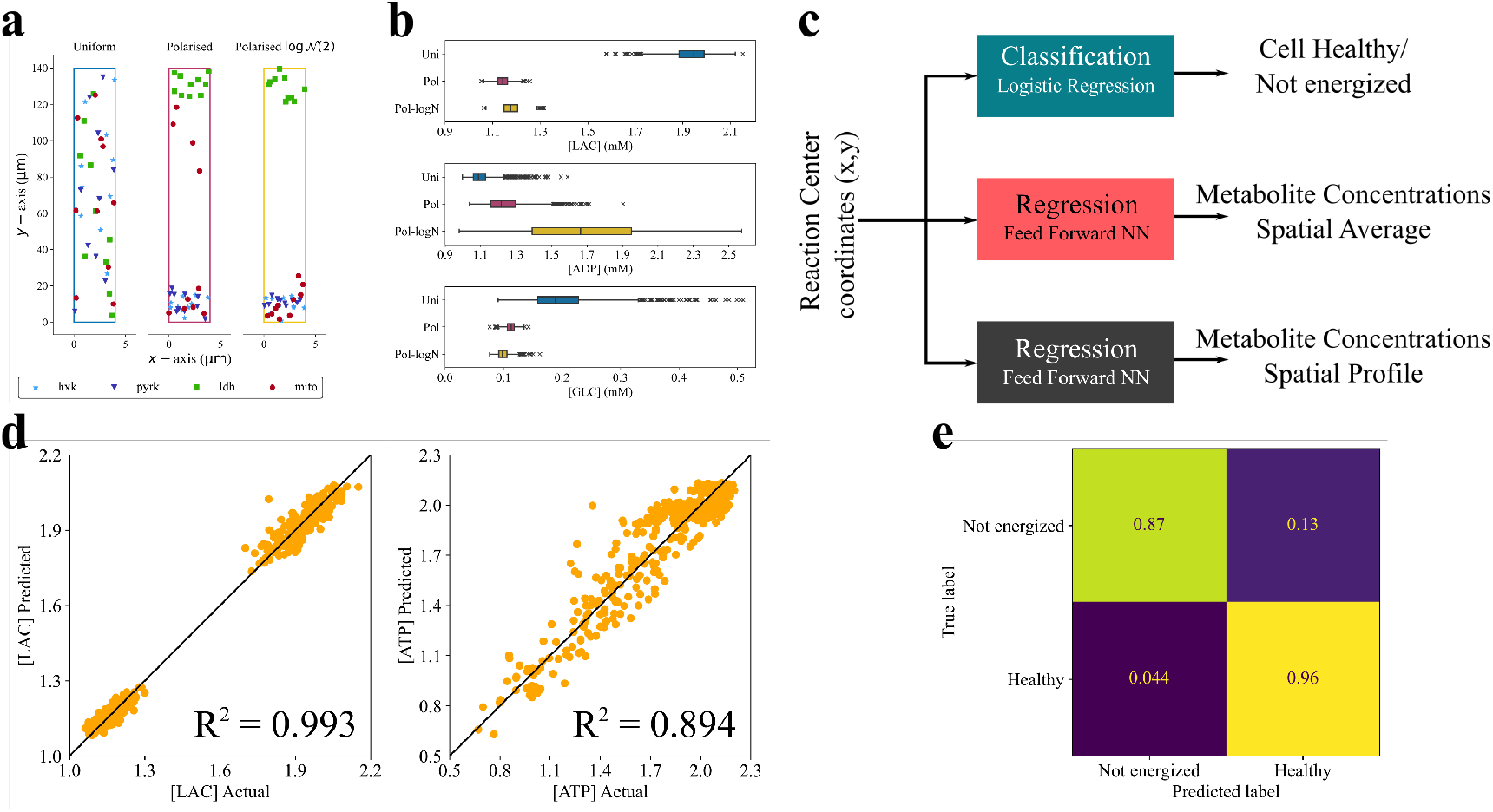
(a) Samples of the three different types of reaction center configurations: uniform, lognormal and polarised lognormal. Glucose enters the cellular domain from the origin of the axis, while lactate exits from the opposite vertex. (b) Distributions of three metabolites of interests for each type of reaction center configuration. (c) The three implemented supervised learning approaches. (d) Out of sample parity plots for the spatial average prediction of [LAC] and [ATP]. (e) Out of sample confusion matrix for the prediction of the energetic state of the cell (non-energized vs healthy).

The dataset used in this study is composed of 1,428 uniform, 1,336 polarized, and 1,314 polarized log-normal realizations. The information for each realization are the *x* and *y* location of the reaction center sites, the average concentration of the six metabolites in the domain at steady state and the spatial concentration at each grid point inside the discretized domain for the six metabolites at the steady state.

### Unsupervised learning

Unsupervised learning algorithms process unlabeled data to discover interesting patterns within the data. For instance, they might perform an association rule analysis or create clusters of similar observations in a dataset (***Hastie et al., 2009b***). Furthermore, dimensionality reduction techniques such as the widely used Principal Component Analysis (PCA), autoencoders (***Wang et al., 2016***), and diffusion maps (***Koronaki et al., 2020, 2023, 2023***) —also fall under the umbrella of unsupervised learning since they provide a reduced data representation without considering the corresponding response variable (or label) of the data.

In the upcoming sections, we will provide a concise overview of the clustering and dimensionality reduction techniques that have been implemented.

#### Clustering

For our clustering study, which aims to discover groups of cells that demonstrate similar metabolic profiles, we implement an *agglomerative* hierarchical clustering algorithm (***Murtagh and Contreras, 2012***; ***Vijaya et al., 2019***). Agglomerative clustering starts with a number of clusters equal to the number of observations and progressively merges clusters until a single cluster remains. The way these clusters merge is based on the dissimilarity metric and the linkage criterion used. Here, the Euclidean distance is implemented as the dissimilarity metric and a ward linkage criterion (***Ward, 1963***). This criterion minimizes the total variance within the cluster by merging the clusters in a way that leads to the smallest increase in variance after each merge. Specifically, it aims to minimize the sum of squared differences within all clusters. The scikit-learn AgglomerativeClustering module is used for this task (***Pedregosa et al., 2011***). Agglomerative hierarchical clustering is selected because it provides insight on how the data merges as the number of clusters changes. This information is readily available in the form of a dendrogram, such as the one presented in Fig. 3.

**Figure 3.**
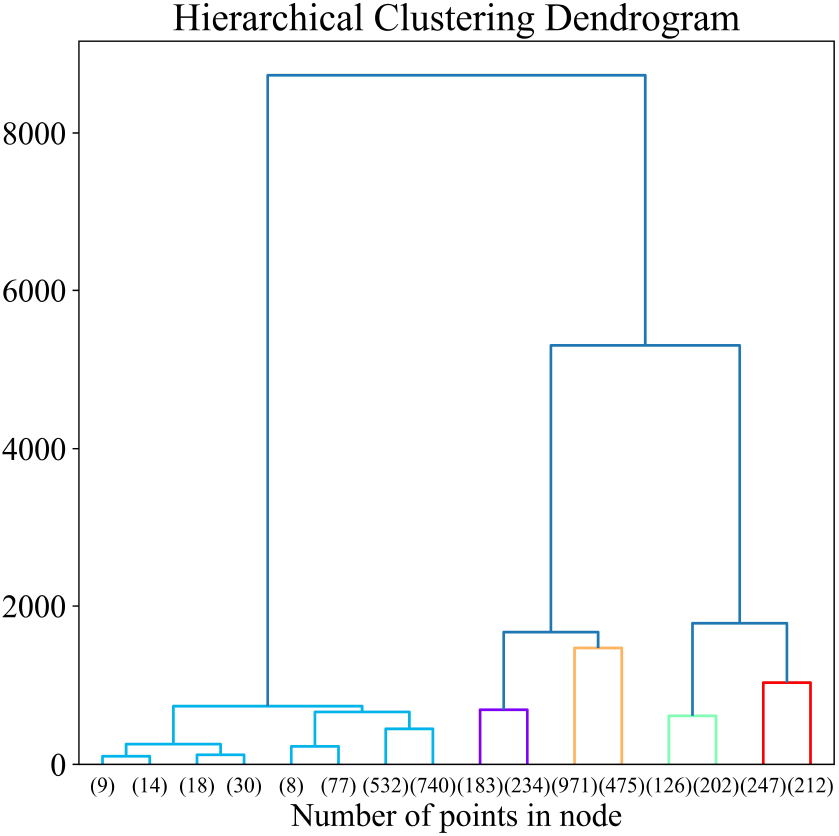
Agglomerative hierarchical clustering dendrogram.

#### Dimensionality reduction

Each row spatial concentration data matrix realization has 79,242 columns, one for each of the six metabolite concentration values at each of the 13,207 grid points. Reducing the dimensionality of the spatial data will be very beneficial when manipulating the data and when training predictive models later on. For this task, Principal Component Analysis (PCA) is implemented (***Jolliffe and Cadima, 2016***). PCA linearly transforms the original data onto a new coordinate system where PCs can be easily identified. The amount of PC retained for subsequent analysis depends on user criteria. In this work, we aim to retain 99.9% of the variance and to keep the reconstruction Root Mean Square Error (RMSE) below 0.03. The scikit-learn PCA module is used for this task (***Pedregosa et al., 2011***).

### Supervised learning

Supervised learning algorithms, in contrast to unsupervised ones, are based on labeled data. In labeled data, the features (*x*_*i*_) are associated with the corresponding responses (*y*_*i*_). These models use available data to make predictions for future observations. Supervised learning encompasses regression for continuous variables and classification for binary or ordinal responses (***James et al., 2021***).

In the present work, the coordinates of the 40 metabolite reaction centers can be considered as inputs, with the response variables being the spatial metabolite concentrations or the spatial average metabolite concentrations. Based on these continuous variables, binary variables can also be engineered (healthy vs. non-energized cell) for classification purposes.

The methods evaluated for this work include: (a) linear methods: for regression, lasso (***Tibshirani, 1996***), and ridge (***Hoerl and Kennard, 1970***) regression, and logistic regression for classification tasks. (b) Support vector machines (SVMs) (***Cortes and Vapnik, 1995***) that can be categorized as linear or nonlinear methods based on the kernel used for classification tasks. (c) Tree-based methods: involving classification and regression trees (***Breiman et al., 1984***) and their ensemble counterparts such as random forests (***Breiman, 2001***), gradient-boosted trees (***Friedman, 2001***), extra trees (***Geurts et al., 2006***), and XGBoost (***Chen and Guestrin, 2016***), which combine numerous trees to improve performance (***Hastie et al., 2009a***). (d) Artificial neural networks (ANN), whose diverse architectures (***Aggarwal, 2018***) can provide valuable options for both classification and regression tasks.

In the present work, logistic regression, random forests, SVM, extra trees, gradient-boosted trees, XGBoost, and ANNs are implemented for supervised learning tasks. However, results are presented only for methods that demonstrate the best performance for our dataset, namely logistic regression for classification tasks and ANNs for regression tasks. An overview of the supervised learning approaches used in this work is presented in Fig. 2c.

Logistic regression is implemented for the classification of cells as healthy or non-energized, based solely on the coordinates of the metabolite reaction centers. The scikit-learn LogisticRegressionCV module is used for this task (***Pedregosa et al., 2011***). This module also has the added benefit of including cross-validation and hyperparameter optimization in the training process, thus reducing overfitting.

In terms of the regression tasks of this work, Artificial Neural Networks (ANNs) are implemented for the prediction of spatial concentration profiles and the spatial average concentrations of the metabolites. The TensorFlow (***Martín Abadi et al., 2015***) and Keras Python libraries (***Chollet et al., 2015***) are used for the development and training of the ANN models in this work.

As the performance of ANNs is significantly influenced by their architecture, optimizing the architecture during the training process is crucial. To achieve this, a Bayesian optimization approach, based on the work of (***Snoek et al., 2012***) is employed for hyperparameter tuning in each ANN model. For this task, we use the keras-tuner Python library (***O’Malley et al., 2019***). Similarly to other optimization methods, Bayesian optimization aims to find optimal values for bounded parameters (hyperparameters in our case), denoted *x*_1_, *x*_2_, …, *x*_*n*_ ∈ *X* that minimize an objective function *f* (*X*) (equivalent to the loss function of the neural network). In Bayesian optimization, a probabilistic model is constructed for *f* (*X*), which allows us to identify the best points in X for evaluating *f* (*X*) in subsequent steps. Unlike local gradient-based methods, this approach considers all available information about *f* (*X*) (***Snoek et al., 2012***).

### SHAP analysis

Shapley values, originally introduced by ***Shapley*** (***1952***) in the field of game theory and proposed as a tool to analyze machine learning models in (***Lundberg and Lee, 2017***), intricately assess the average contribution of each feature’s value to predictions. In this way, they provide an understanding of how perturbations of a variable can influence the output of the model, thus shedding light on models that have traditionally been considered “black boxes”.

In this work, a SHAP analysis is performed on 3 of the ANN regression models developed for the prediction of spatial average metabolite concentrations. The final goal is to discover not only the inputs have the most influence on model output, but also the type of influence they have. However, the results of this SHAP analysis only provide information about the relationships between inputs and *model outputs*. These results shed light on previously “opaque” ML methods but should not be used to make causal claims about input/output relationships.

## Results

### Clustering

An agglomerative hierarchical clustering algorithm is used to discover observations with similar characteristics solely based on the results (outputs) of the computational model. The results consist of the concentration values of the six metabolites at the 13,207 grid points used for the computational simulations. A subset of the entire dataset is used for clustering. Subsequently, the input to the clustering algorithm is a matrix of dimension 1,767×79,242.

Following clustering, we attempt to analyze the resulting realization groups. We investigate a) the mean spatial averages of the six different metabolite concentrations, b) the mean [ATP] ∶ [ADP], and c) the distribution of the *y* coordinates of mitochondria in each cluster. The mean spatial averages of all six metabolites provide a great overview of the metabolic state of the cell. The mean [ATP] ∶ [ADP] is an excellent indicator of cell health, as cells that demonstrate a ratio lower than 1 can be considered non-energized and in a state of deterioration, while cells with a ratio greater than 1 are considered adequately energized and thus healthy. Finally, by investigating the distribution of the *y*-axis coordinates of the mitochondria for each cluster, we can uncover possible relationships between the metabolic state of the cell and the locations of the mitochondria. Our analysis reveals the following.

1. Based on the results of Table 1, it appears that cluster 1 contains healthier cells, given the fact that the average [ATP] ∶ [ADP] is the highest of the 5 clusters with a value of 1.910. It also appears that cluster 1 has the lowest values for [GLY] and [PYR], suggesting that this is the most efficient cluster that consumes these two substrates to maximize ATP production. Cluster 3 contains fewer energized cells, given its average [ATP] ∶ [ADP] of 0.435. It should also be noted that cluster 1 contains only realizations with a uniform reaction site distribution. This is also evident in Fig. 5d.
2. When comparing the clusters containing realizations of non-uniform reaction center distributions, we can see that for clusters 2 and 3 the mitochondria are located close to the cell inlet (and subsequently closer to the glucose entering the cell), whereas the cells for clusters 0 and 4, the mitochondria have better coverage of the spatial domain. This result is visualized in the histogram of Fig. 5d. This is a hint that mitochondrial distribution is a great driver of cell health, as clusters 2 and 3 demonstrate a lower average [ATP] ∶ [ADP] (0.852 and 0.435, respectively) when compared to clusters 0 and 4 (1.610 and 1.388, respectively).
3. Last, solely based on the concentration values the algorithm can discern the three main groups (uniform, polarized and polarized log-normal) of cells available in the dataset. Cluster 1 contains solely uniform realizations, clusters 0 and 4 contain mostly polarized and some polarized log-normal realizations. Clusters 2 and 3 contain almost exclusively polarized lognormal realizations.

**Table 1.**
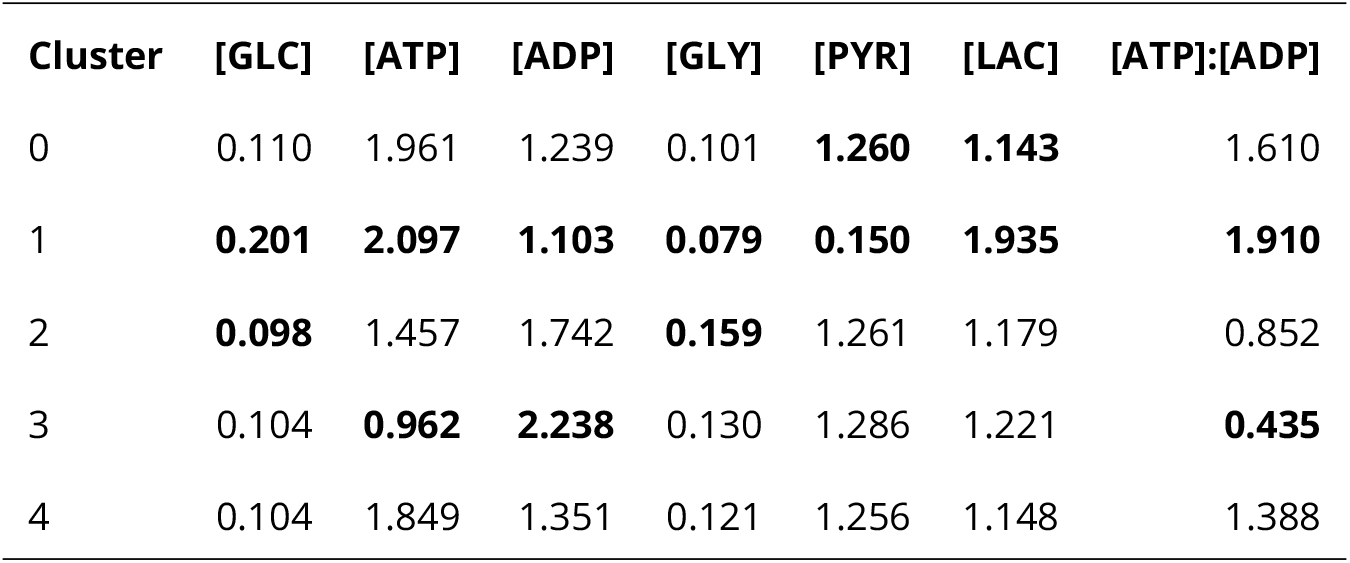
Mean spatial metabolite concentration averages in each cluster. Lowest and highest values for each metabolite are presented in bold.

| Cluster | [GLC] | [ATP] | [ADP] | [GLY] | [PYR] | [LAC] | [ATP]:[ADP] |
| --- | --- | --- | --- | --- | --- | --- | --- |
| 0 | 0.110 | 1.961 | 1.239 | 0.101 | **1.260** | **1.143** | 1.610 |
| 1 | **0.201** | **2.097** | **1.103** | **0.079** | **0.150** | **1.935** | **1.910** |
| 2 | **0.098** | 1.457 | 1.742 | **0.159** | 1.261 | 1.179 | 0.852 |
| 3 | 0.104 | **0.962** | **2.238** | 0.130 | 1.286 | 1.221 | **0.435** |
| 4 | 0.104 | 1.849 | 1.351 | 0.121 | 1.256 | 1.148 | 1.388 |

### Discerning between Healthy and Not energized cells

As already established, an important indicator of cell health is the spatial average of [ATP] ∶ [ADP] (ATP-to-ADP ratio) within the cell. When [ATP] ∶ [ADP] *≥* 1, the cell is considered adequately energized and healthy, whereas when [ATP] ∶ [ADP] < 1 the cell is considered non-energized and unhealthy.

Given this threshold and the calculated spatial averages of [ATP] ∶ [ADP] for all available samples, we can convert the continuous output (ratio) to binary (health), where *health* = 0 when *ratio* < 1, and *health* = 1 when *ratio ≥* 1. Using the available reaction center coordinates for each observation as inputs and the binary variable *health* as output, we can train a logistic regression model that can predict the health status of a cell, given only the coordinates of the reaction centers.

The inputs consist of the coordinates (*x, y*) for 40 different reaction centers (10 of each of the four types), totaling 80 inputs. Before training the logistic regression model, the inputs are centered to 0 and scaled by their standard deviation.

Of the 4,078 observations, 85% is used as a training set and 15% as a test set. Checking the performance of the model on both sets can allow us to avoid overfitting and make sure that the resulting model generalizes well in unseen data.

The resulting logistic regression model can discern between healthy and non-energized cells with high accuracy for both the training (0.9412) and the test set (0.9515). Further classification metrics, such as the f1 score, recall, and precision, are presented in Table 2. The confusion matrices for the performance of the logistic regression algorithm in both the training and the test set are presented in Fig. 2e.

**Table 2.** Binary classification metrics. The trained logistic regression model shows great performance for both the test and training sets.

|  | Training Set | Test Set |
| --- | --- | --- |
| F1 | 0.9638 | 0.9701 |
| Accuracy | 0.9412 | 0.9515 |
| Precision | 0.9561 | 0.9650 |
| Recall | 0.9716 | 0.9753 |

Given the nature of the logistic regression algorithm, we can use the resulting coefficients to try to make sense of the connection between the input variables and the output (cell health state). Observing Fig. 4, it is evident that the y coordinates of the mitochondria highly influence the model output.

**Figure 4.**
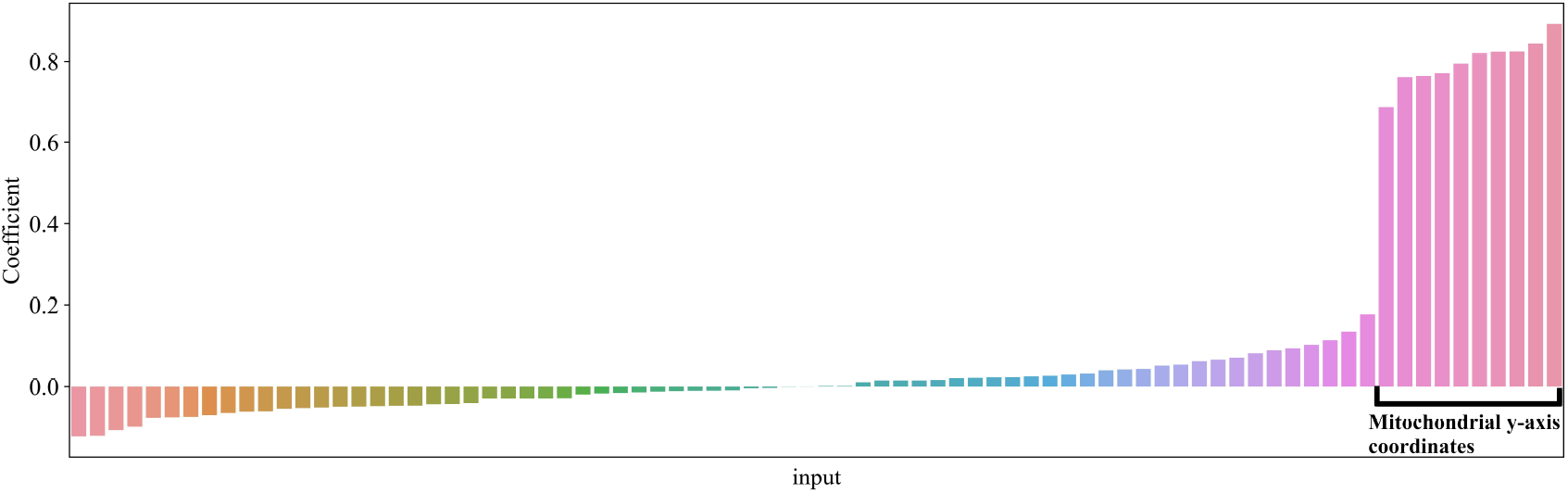
Logistic regression coefficients for the not energized/ healthy cell classification problem. The y-axis coordinates of the mitochondria appear to be highly influential for the model. Essentially, the dominating coefficient values of the mitochondrial y-axis coordinates show that a better spread of mitochondria within the cell -and not close to the cell inlet-increases the probability of a healthy cell.

**Figure 5.**
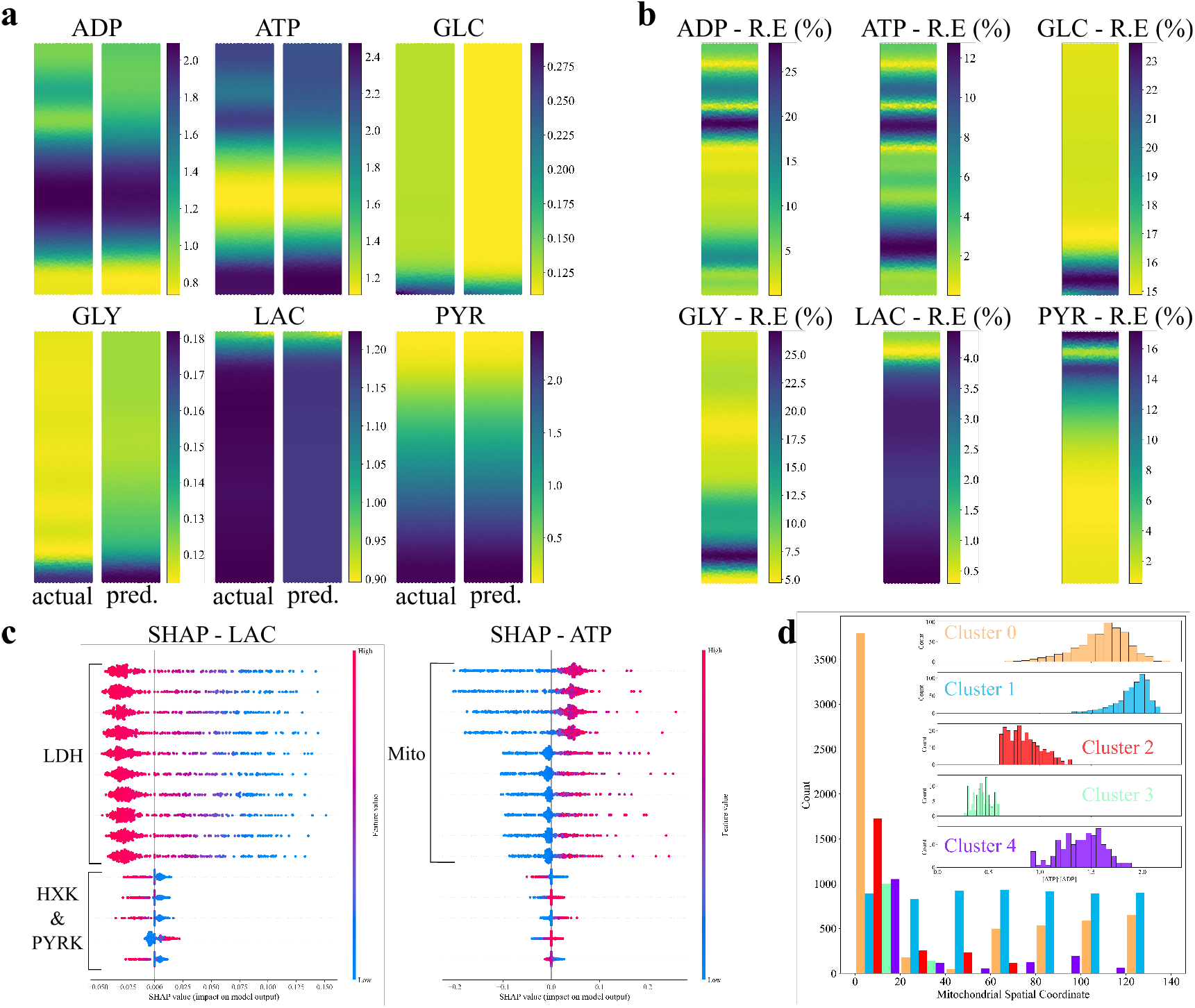
(a) Actual and predicted spatial metabolite concentration (in mM) profiles for randomly sampled, out of sample reaction center configurations. (b) Absolute relative error (%) between the actual and predicted metabolite concentrations presented in (a). (c) SHAP values for the spatial average [LAC] and [ADP] predictive models. For the [LAC] model, the location of the LDH reaction centers is the most important for the output of the model, followed by the locations of HXK and PYRK reaction centers. For the [ATP] model, the location of the mitochondrial reaction centers is the most important for the model output. (d) Histogram showing the distribution of the mitochondrial reaction center for each discovered cluster. On the top right, histograms of the [ATP] ∶ [ADP] for each cluster. Clusters 0, 1 and 4 appear to have more observations with [ATP] ∶ [ADP]>1, whereas clusters 2 and 3 appear to be more problematic. When investigating the y coordinates of the mitochondria for each cluster, it is clear that for clusters 2 and 3, the mitochondria are located primarily close to the input of the cell (*y*_Mito_ = 0), while for clusters 0, 1 and 4 the mitochondria have better coverage of the domain.

### Predicting spatial metabolite concentrations

Although the prediction of cell health status is already quite valuable, it is often preferable to have models that can predict a continuous variable rather than a binary variable. Similarly to the classification problem, the coordinates of the 40 reaction centers can be used to predict continuous variables. These continuous values can be either the spatial averages of the six metabolite concentrations or, taking it one step further, the spatial profiles of the metabolite concentrations.

As in the binary classification problem, 85% of the 4,078 observations are used as a training set and 15% as a test set.

### Spatial averages

ANN models are used for the prediction of spatial averages of the six metabolite concentrations. A different model is trained for each metabolite concentration. To ensure optimal performance, we include hyperparameter optimization in model training. The hyperparameters tested are:

1. The number of network layers (*N*_layers_). Varied between 2 and 5.
2. The number of neurons per layer (*N*_neurons_). Varied between 2 and 100.
3. The activation function used in all of the layers. The functions tested are: a) sigmoid, b) tanh, and c) Relu. It should be noted that the output layer always has a single neuron using a linear activation function.

50% of the test set is used as a validation set during training to avoid overfitting through early stopping. This means that if the validation error does not drop after a certain number of training epochs (10 in our case), the training stops.

The resulting neural networks, along with their performance in the test set, are presented in Table 3. The models developed for the prediction of [PYR] and [LAC] demonstrate excellent performance with a R^2^ = 0.998 and R^2^ = 0.993 respectively. Satisfactory performance is observed for the predictions of [ADP] (R^2^ = 0.914), [ATP] (R^2^ = 0.894) and [GLC] R^2^ = 0.862). The developed [GLY] regression model shows a rather modest performance (R^2^ = 0.595).

**Table 3.**
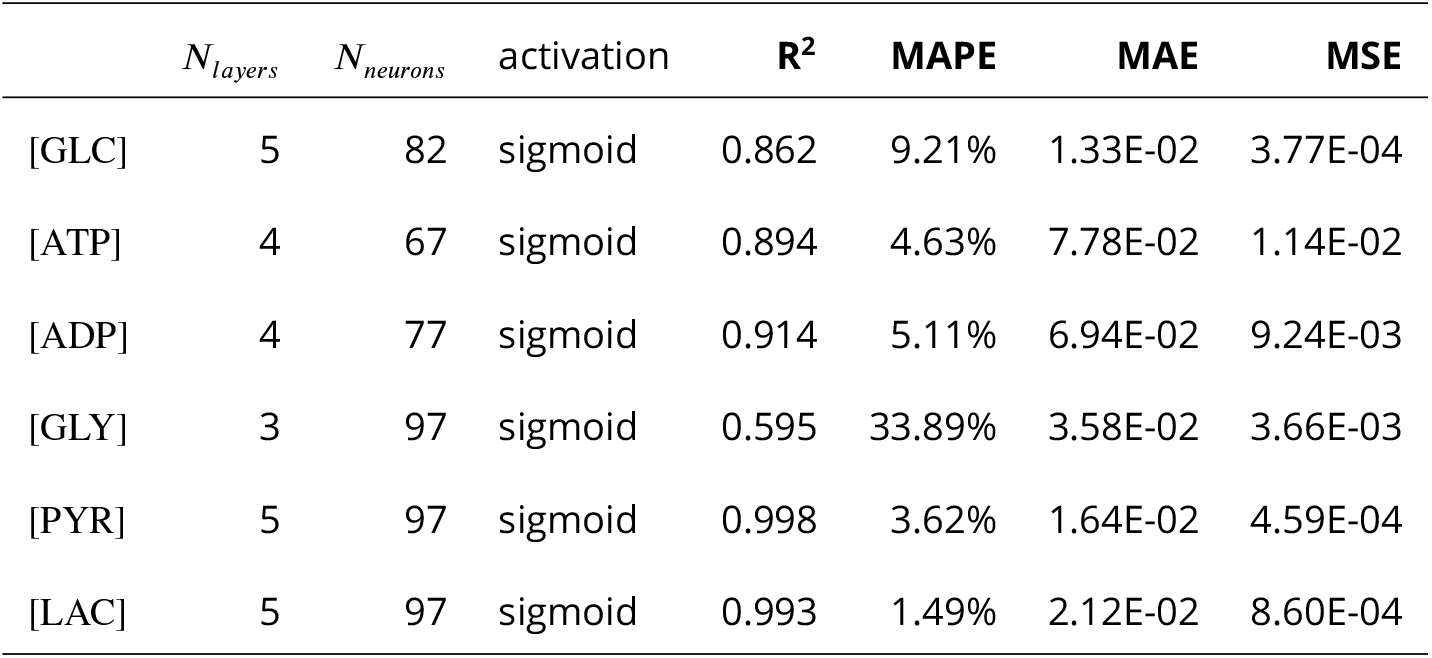
Optimal hyperparameters and spatial average metabolite concentration regression test set performance metrics for the developed ANN models. Results for [PYR] and [LAC] are excellent. For [GLC], [ATP] and [ADP], the results are satisfactory. However, the [GLY] model demonstrates a slightly worse performance.

| | $N_{layers}$ | $N_{neurons}$ | activation | $R^2$ | MAPE | MAE | MSE |
| --- | --- | --- | --- | --- | --- | --- | --- |
| [GLC] | 5 | 82 | sigmoid | 0.862 | 9.21% | 1.33E-02 | 3.77E-04 |
| [ATP] | 4 | 67 | sigmoid | 0.894 | 4.63% | 7.78E-02 | 1.14E-02 |
| [ADP] | 4 | 77 | sigmoid | 0.914 | 5.11% | 6.94E-02 | 9.24E-03 |
| [GLY] | 3 | 97 | sigmoid | 0.595 | 33.89% | 3.58E-02 | 3.66E-03 |
| [PYR] | 5 | 97 | sigmoid | 0.998 | 3.62% | 1.64E-02 | 4.59E-04 |
| [LAC] | 5 | 97 | sigmoid | 0.993 | 1.49% | 2.12E-02 | 8.60E-04 |

### Spatial profiles

Given the satisfactory performance of the predictive models for spatial average concentrations, we can go a step further, trying to predict the spatial concentration profiles of the six metabolites in the cells given only the positions of the 40 reaction centers. Given the nature of the output vector (length of 79,242 -six metabolite concentration values for each of the 13,207 grid points), it is clear that dimensionality reduction methods can be useful for reducing the dimensions of the output space.

A model is developed for the spatial concentration profile of each metabolite. This translates to an output vector of length 13,207 (1 concentration for each grid point) for each developed model.

Principal component analysis (PCA) is used for this task, following the centering of the output data using their mean and their scaling using their standard deviation. The number of principal components retained for each output is chosen based on the reconstruction root mean squared error (RMSE). The number of principal components retained is chosen so that the reconstruction root mean squared error (RMSE) is lower than 0.03. This leads to a different number of principal components (PCs) for each metabolite. Specifically, 17 PCs are retained for the spatial [ATP] and spatial [ADP] models, 5 PCs are kept for the spatial [PYR] and [LAC] models, 9 PCs are retained for the spatial [GLY] model, and finally, 11 PCs are used for the spatial [GLC] model.

As with the spatial average concentration predictive models, we include hyperparameter optimization when training the spatial profile models. The hyperparameters tested are:

1. The number of network layers (*N*_layers_). Varied between 2 and 5.
2. The number of neurons per layer (*N*_neurons_). Varied between 10 and 650.
3. The activation function used in all of the layers. The functions tested are a) sigmoid, b) tanh, c) relu, and d) elu. Note that the output layer size is equal to the number of PCs retained for the model. Furthermore, output neurons always use a linear activation function.

Once again, 50% of the test set is used as a validation set and early stopping is used during training to avoid overfitting.

The resulting neural networks, along with their performance in the test set, are presented in Table 4. The models developed for the spatial concentration prediction of [PYR] and [LAC] demonstrate excellent performance with R^2^ = 0.956 and R^2^ = 0.972 respectively. Reasonable performance is observed for the predictions of [ADP] (R^2^ = 0.698), [ATP] (R^2^ = 0.700) and [GLC] R^2^ = 0.775). The developed [GLY] regression model shows unsatisfactory performance (R^2^ = 0.474). It is evident that the performance of the models follows the trend of the models trained for simpler spatial average concentration predictions.

**Table 4.** Optimal hyperparameters and spatial profile metabolite concentration regression test set performance metrics for the developed ANN models. Performance for [PYR] and [LAC] are excellent. For [GLC], [ATP] and [ADP], the results are reasonable. However, the [GLY] model demonstrates a non-satisfactory performance.

| | $N_{layers}$ | $N_{neurons}$ | activation | $R^2$ | MAPE | MAE | MSE |
| --- | --- | --- | --- | --- | --- | --- | --- |
| [GLC] | 2 | 285 | sigmoid | 0.775 | 9.24% | 1.34E-02 | 6.16E-04 |
| [ATP] | 3 | 500 | sigmoid | 0.700 | 12.93% | 1.69E-01 | 6.31E-02 |
| [ADP] | 3 | 650 | sigmoid | 0.698 | 13.10% | 1.69E-01 | 6.24E-02 |
| [GLY] | 5 | 340 | sigmoid | 0.474 | 64.43% | 4.40E-02 | 5.78E-03 |
| [PYR] | 5 | 10 | elu | 0.956 | 9.33% | 2.42E-02 | 1.47E-03 |
| [LAC] | 5 | 10 | elu | 0.972 | 1.40% | 2.17E-02 | 1.43E-03 |

Some examples of predicted spatial metabolite concentrations are presented in Fig. 5, along with the actual metabolite concentration profile and the absolute relative reconstruction error (%). It is evident that, while most of the trends are retained, there are certain zones of higher relative errors for certain models. However, it should be noted that for certain metabolite concentrations, the actual values can be quite small, which means that small absolute errors in the prediction can lead to very high relative errors.

### Effect of inputs on model output (SHAP analysis)

Following the training of individual models for both spatial metabolite concentrations and spatial average concentrations, we would like to investigate the influence of model inputs on model outputs.

For this task, we will focus on three spatial average concentration models. Specifically, we will perform a SHAP analysis for the spatial average predictive models of [LAC], [ATP], and [ADP]. The results of the SHAP analysis for [LAC] and [ATP] are presented in Fig. 5c.

For the [LAC] model, it appears that the positions of the LDH reaction centers on the y-axis are the most influential variables for the model output. Furthermore, it appears that when the reaction centers are located high on the y-axis, the model output is negatively influenced.

For the [ATP] model, it appears that the positions of the mitochondria on the y-axis are the most influential variables for the model output. Furthermore, we can observe that when the mitochondria are located close to the inlet of the cell, the predicted spatial average [ATP] is negatively affected.

For the [ADP] model, it appears that the positions of the mitochondria on the y-axis are the most influential variables for the model output. Furthermore, we can observe that when the mito-chondria are located close to the outlet of the cell, the predicted spatial average [ADP] is negatively affected.

The results of the SHAP analysis for the [ATP] and [ADP] models appear to be mirror images of each other. It looks like when the mitochondria are located close to the cell inlet, the average [ATP] of the cell is negatively affected, while the average [ADP] is positively affected. However, when the mitochondria are located close to the cell outlet, the average [ATP] appears to be positively affected, while the average [ADP] is negatively affected.

It is worth pointing out that although SHAP analysis can shed light on the relationship between model inputs and outputs, it cannot be used to make causal claims regarding these relationships.

## Conclusions & Perspectives

Several machine learning methods are implemented to gain insight into metabolism within astrocyte cells by exploring a dataset consisting of static metabolic images of astrocytes generated from a reaction-diffusion computational model solved in a two-dimensional geometrical domain.

Hierarchical clustering of the dataset, based solely on the outputs of the computational experiments, results in 5 groups. Each group has its distinctive metabolic characteristics. The discovered clusters reveal differences not only regarding the average metabolite concentrations of their members but also on the side of the inputs (reaction centers). By analyzing the reaction center configurations corresponding to each cluster, we observe that a uniform mitochondrial distribution results in the best overall cell health. On the contrary, realizations of polarized log-normal mitochondrial distributions exhibit the worst overall cell health. It is also shown that a high concentration of mitochondria close to the cell inlet is not beneficial to the cell, especially compared to a better coverage of the cell. Our results are in agreement with the discussion presented on the original model ***Farina et al***. (***2023***).

Following the encouraging results of the clustering analysis, we developed a classification model able to accurately predict cell health (healthy/non-energized) taking only the reaction center positions as input. The nature of the logistic regression model used also allows us to confirm the importance of mitochondrial distribution on cell health.

Moving one step forward and past the binary nature of healthy/non-energized output, we constructed artificial neural network (ANN) regression models that can satisfactorily predict the average spatial concentration of the six metabolites of interest. We show that it is also possible to recreate the spatial profiles for the six metabolites with satisfactory accuracy. For both regression problems, the LAC and PYR predictive models perform best, followed by those for ATP, ADP, and GLC. The GLY models exhibit only a modest performance. In general, regression models that predict spatial average metabolite concentrations perform better than their counterparts that predict the *full* spatial profiles.

Following regression, SHAP analysis allows us to see into the black boxes that are ANNs, high-lighting the importance and influence of different reaction center positions on model output. Specifically, the positions of the lactate dehydrogenase (LDH) reaction centers greatly influence the spatial average concentration of [LAC], and the positions of the mitochondria influence the spatial average concentration of [ATP] and [ADP], consistent with the formulation of the metabolic model.

Based on these encouraging results, several future steps are possible. The first step would involve applying a similar approach to more complex computational domains that better represent actual astrocytic cells. Going further, we could use existing experimental images that provide mito-chondrial location data ***Chu et al***. (***2022***) as initial inputs for our approach, with a suitable choice of computational model. Extending this methodology to different or more detailed metabolic pathways could also provide insights into a broader range of species that influence metabolism.

While in this paper we have considered a simplified two-dimensional domain and a basic metabolic model, our machine learning approach can serve as a bridge linking the spatial information from *in vitro* or *in vivo* images with computational models, thereby gaining deeper insights into metabolic dynamics and cellular states. It should be highlighted that the methods used in this work are not limited to the dataset presented; they can be readily applied to other datasets derived from various metabolic computational models.

## Acknowledgments

The authors thank Dr. Michela Bernini for the sketch of Figure 1a. PP gratefully acknowledges funding from the FSTM in the University of Luxembourg.

## References

Agapakis CM, Boyle PM, Silver PA. Natural strategies for the spatial optimization of metabolism in synthetic biology. Nature chemical biology. 2012; 8(6):527–535.

Aggarwal CC. Neural Networks and Deep Learning: A Textbook. Cham: Springer International Publishing; 2018. doi: 10.1007/978-3-319-94463-0.

Alber M, Buganza Tepole A, Cannon WR, D. S, Dura-Bernal S, Garikipati K, Karniadakis G, Lytton WW, Perdikaris P, Petzold L, et al. Integrating machine learning and multiscale modeling—perspectives, challenges, and opportunities in the biological, biomedical, and behavioral sciences. NPJ digital medicine. 2019; 2(1):115.

Alexandrov T. Spatial metabolomics: from a niche field towards a driver of innovation. Nature Metabolism. 2023; 5(9):1443–1445.

Alnæs M, Blechta J, Hake J, Johansson A, Kehlet B, Logg A, Richardson C, Ring J, Rognes ME, Wells GN. The FEniCS project version 1.5. Archive of numerical software. 2015; 3(100).

Aubert A, Costalat R. Interaction between astrocytes and neurons studied using a mathematical model of compartmentalized energy metabolism. Journal of Cerebral Blood Flow & Metabolism. 2005; 25(11):1476– 1490.

Aubert A, Costalat R, Magistretti PJ, Pellerin L. Brain lactate kinetics: modeling evidence for neuronal lactate uptake upon activation. Proceedings of the National Academy of Sciences. 2005; 102(45):16448–16453.

Bell M, Bartol T, Sejnowski T, Rangamani P. Dendritic spine geometry and spine apparatus organization govern the spatiotemporal dynamics of calcium. Journal of General Physiology. 2019; 151(8):1017–1034.

Breiman L. Random Forests. Machine Learning. 2001 Oct; 45(1):5–32. doi: 10.1023/A:1010933404324.

Breiman L, Friedman J, Olshen RA, Stone CJ. Classification and Regression Trees. New York: Chapman and Hall/CRC; 1984. doi: 10.1201/9781315139470.

Chen T, Guestrin C. XGBoost: A Scalable Tree Boosting System. In: Proceedings of the 22nd ACM SIGKDD International Conference on Knowledge Discovery and Data Mining San Francisco California USA: ACM; 2016. p. 785–794. doi: 10.1145/2939672.2939785.

Chollet F, et al., Keras; 2015. https://keras.io.

Chu CH, Tseng WW, Hsu CM, Wei AC. Image analysis of the mitochondrial network morphology with applications in cancer research. Frontiers in Physics. 2022; 10:855775.

Cleri F. Agent-based model of multicellular tumor spheroid evolution including cell metabolism. The European Physical Journal E. 2019; 42:1–15.

Cortassa S, Aon MA, Marbán E, Winslow RL, O’Rourke B. An integrated model of cardiac mitochondrial energy metabolism and calcium dynamics. Biophysical journal. 2003; 84(4):2734–2755.

Cortes C, Vapnik V. Support-Vector Networks. Machine Learning. 1995 Sep; 20(3):273–297. doi: 10.1007/BF00994018.

Ellingsrud AJ, Solbrå A, Einevoll GT, Halnes G, Rognes ME. Finite element simulation of ionic electrodiffusion in cellular geometries. Frontiers in Neuroinformatics. 2020; 14:11.

Farina S, Claus S, Hale JS, Skupin A, Bordas SP. A cut finite element method for spatially resolved energy metabolism models in complex neuro-cell morphologies with minimal remeshing. Advanced Modeling and Simulation in Engineering Sciences. 2021; 8(1):1–32.

Farina S, Voorsluijs V, Fixemer S, Bouvier D, Claus S, Ellisman M, Bordas SP, Skupin A. Mechanistic multiscale modelling of energy metabolism in human astrocytes reveals the impact of morphology changes in Alzheimer’s Disease. PLOS Computational Biology. 2023; 19(9):e1011464.

Friedman JH. Greedy Function Approximation: A Gradient Boosting Machine. The Annals of Statistics. 2001; 29(5):1189–1232.

Garcia GC, Gupta K, Bartol TM, Sejnowski TJ, Rangamani P. Mitochondrial morphology governs ATP production rate. Journal of General Physiology. 2023; 155(9):e202213263.

Geurts P, Ernst D, Wehenkel L. Extremely Randomized Trees. Machine Learning. 2006 Apr; 63(1):3–42. doi: 10.1007/s10994-006-6226-1.

Hastie T, Tibshirani R, Friedman J. Ensemble Learning. In: Hastie T, Tibshirani R, Friedman J, editors. The Elements of Statistical Learning: Data Mining, Inference, and Prediction New York, NY: Springer; 2009. p. 605–624. doi: 10.1007/978-0-387-84858-7_16.

Hastie T, Tibshirani R, Friedman J. Unsupervised Learning. In: Hastie T, Tibshirani R, Friedman J, editors. The Elements of Statistical Learning: Data Mining, Inference, and Prediction Springer Series in Statistics, New York, NY: Springer; 2009. p. 485–585. doi: 10.1007/978-0-387-84858-7_14.

Hoerl AE, Kennard RW. Ridge Regression: Biased Estimation for Nonorthogonal Problems. Technometrics. 1970 Feb; 12(1):55–67. doi: 10.1080/00401706.1970.10488634.

Hughes TJ. The finite element method: linear static and dynamic finite element analysis. Courier Corporation; 2012.

Jackson JG, Robinson MB. Reciprocal regulation of mitochondrial dynamics and calcium signaling in astrocyte processes. Journal of Neuroscience. 2015; 35(45):15199–15213.

James G, Witten D, Hastie T, Tibshirani R. In: Statistical Learning New York, NY: Springer US; 2021. p. 15–57. doi: 10.1007/978-1-0716-1418-1_2.

Jolliffe IT, Cadima J. Principal component analysis: a review and recent developments. Philosophical transactions of the royal society A: Mathematical, Physical and Engineering Sciences. 2016; 374(2065):20150202.

Khalid MU, Tervonen A, Korkka I, Hyttinen J, Lenk K. Geometry-based computational modeling of calcium signaling in an astrocyte. In: EMBEC & NBC 2017: Joint Conference of the European Medical and Biological Engineering Conference (EMBEC) and the Nordic-Baltic Conference on Biomedical Engineering and Medical Physics (NBC), Tampere, Finland, June 2017 Springer; 2018. p. 157–160.

Kitano H. Computational systems biology. Nature. 2002; 420(6912):206–210.

Kobayashi M, Takeda M, Sato T, Yamazaki Y, Kaneko K, Ito KI, Kato H, Inaba H. In vivo imaging of spontaneous ultraweak photon emission from a rat’s brain correlated with cerebral energy metabolism and oxidative stress. Neuroscience research. 1999; 34(2):103–113.

Koronaki ED, Nikas AM, Boudouvis AG. A Data-Driven Reduced-Order Model of Nonlinear Processes Based on Diffusion Maps and Artificial Neural Networks. Chemical Engineering Journal. 2020 Oct; 397:125475. doi: 10.1016/j.cej.2020.125475.

Koronaki ED, Evangelou N, Psarellis YM, Boudouvis AG, Kevrekidis IG. From Partial Data to Out-of-Sample Parameter and Observation Estimation with Diffusion Maps and Geometric Harmonics. Computers & Chemical Engineering. 2023 Jul; p. 108357. doi: 10.1016/j.compchemeng.2023.108357.

Llaneras F, Picó J. Stoichiometric modelling of cell metabolism. Journal of bioscience and bioengineering. 2008; 105(1):1–11.

Lundberg SM, Lee SI. A Unified Approach to Interpreting Model Predictions. In: Advances in Neural Information Processing Systems, vol. 30 Curran Associates, Inc.; 2017.

Martín Abadi, Ashish Agarwal, Paul Barham, Eugene Brevdo, Zhifeng Chen, Craig Citro, Greg S Corrado, Andy Davis, Jeffrey Dean, Matthieu Devin, Sanjay Ghemawat, Ian Goodfellow, Andrew Harp, Geoffrey Irving, Michael Isard, Jia Y, Rafal Jozefowicz, Lukasz Kaiser, Manjunath Kudlur, Josh Levenberg, et al., TensorFlow: Large-scale Machine Learning on Heterogeneous Systems; 2015.

Murray JD. Mathematical Biology: I. An introduction. Springer; 2002.

Murtagh F, Contreras P. Algorithms for Hierarchical Clustering: An Overview. WIREs Data Mining and Knowledge Discovery. 2012; 2(1):86–97. doi: 10.1002/widm.53.

O’Malley T, Bursztein E, Long J, Chollet F, Jin H, Invernizzi L, et al., Keras Tuner; 2019. https://github.com/keras-team/keras-tuner.

Pedregosa F, Varoquaux G, Gramfort A, Michel V, Thirion B, Grisel O, Blondel M, Prettenhofer P, Weiss R, Dubourg V, Vanderplas J, Passos A, Cournapeau D, Brucher M, Perrot M, Duchesnay E. Scikit-learn: Machine Learning in Python. Journal of Machine Learning Research. 2011; 12:2825–2830.

Pellerin L, Pellegri G, Bittar PG, Charnay Y, Bouras C, Martin JL, Stella N, Magistretti PJ. Evidence supporting the existence of an activity-dependent astrocyte-neuron lactate shuttle. Developmental neuroscience. 1998; 20(4-5):291–299.

Quarteroni A, Valli A. Numerical approximation of partial differential equations, vol. 23. Springer Science & Business Media; 2008.

Sánchez-Alvarez R, Tabernero A, Medina JM. Endothelin-1 stimulates the translocation and upregulation of both glucose transporter and hexokinase in astrocytes: relationship with gap junctional communication. Journal of neurochemistry. 2004; 89(3):703–714.

Shapley LS. A Value for N-Person Games. RAND Corporation; 1952.

Snoek J, Larochelle H, Adams RP, Practical Bayesian Optimization of Machine Learning Algorithms; 2012.

Suzuki R, Hotta K, Oka K. Spatiotemporal quantification of subcellular ATP levels in a single HeLa cell during changes in morphology. Scientific reports. 2015; 5(1):16874.

Szabó A, Merks RM. Cellular potts modeling of tumor growth, tumor invasion, and tumor evolution. Frontiers in oncology. 2013; 3:87.

Tantama M, Yellen G. Imaging Changes in the Cytosolic ATP-to-ADP Ratio. Methods in enzymology. 2014; 547:355–371. doi: 10.1016/B978-0-12-801415-8.00017-5.

Tibshirani R. Regression Shrinkage and Selection via the Lasso. Journal of the Royal Statistical Society Series B (Methodological). 1996; 58(1):267–288. doi: 10.1111/j.2517-6161.1996.tb02080.x.

Vijaya, Sharma S, Batra N. Comparative Study of Single Linkage, Complete Linkage, and Ward Method of Agglomerative Clustering. In: 2019 International Conference on Machine Learning, Big Data, Cloud and Parallel Computing (COMITCon) Faridabad, India: IEEE; 2019. p. 568–573. doi: 10.1109/COMITCon.2019.8862232.

Wang Y, Yao H, Zhao S. Auto-Encoder Based Dimensionality Reduction. Neurocomputing. 2016 Apr; 184:232–242. doi: 10.1016/j.neucom.2015.08.104.

Ward JH. Hierarchical Grouping to Optimize an Objective Function. Journal of the American Statistical Association. 1963 Mar; 58(301):236–244. doi: 10.1080/01621459.1963.10500845.

Zampieri G, Vijayakumar S, Yaneske E, Angione C. Machine and deep learning meet genome-scale metabolic modeling. PLoS computational biology. 2019; 15(7):e1007084.

